# Feline Papillomavirus Strain P20 Assembled from Metagenomic Data Isolated from the Human Skin of a House Cat Owner

**DOI:** 10.1101/2021.11.01.466825

**Authors:** Ema H. Graham, Michael S. Adamowicz, Peter C Angeletti, Jennifer L. Clarke, Samodha C. Fernando, Joshua R. Herr

**Author notes:** Corresponding Author, Joshua R. Herr, PhD; 422 Plant Sciences Hall, University of Nebraska, Lincoln NE 68583.

## Abstract

A feline papillomavirus was assembled from metagenomic sequencing data collected from the human skin of a house cat owner. This circular papillomavirus strain P20 is 8069 bp in length, has a GC content of 54.38%, and displays genome organization typical of feline papillomaviruses with six annotated protein coding regions. The genome exhibits approximately 75% sequence similarity to other feline papillomavirus genomes.

## ANNOUNCEMENT

Papillomaviruses are most recognized for infecting humans, but they also broadly infect mammals, reptiles, and birds (1). Classification of papillomaviruses has largely focused on the sequence divergence of the Major Capsid Protein L1. High conservation of the L1 protein across papillomaviruses suggests their evolution mirrors the phylogeny of their hosts (2). Recent studies have reported on the discovery and diversity of papillomaviruses associated with domestic and wild cats (3). Seven types of feline papillomaviruses are currently recognized that may or may not produce both skin and oral squamous cell carcinomas in house cats (4).

In a previous study we addressed the diversity and stability of DNA viruses on human skin by sampling three anatomical locations – left hand, right hand, and scalp – over a longitudinal 6-month period for 43 human subjects (5). Briefly, samples were collected using a tandem dry and wet swab technique. Swabs were then saturated in 1x PBS and were centrifuged at 20,000xg. Sample eluent was then run through a 0.22 μm filter to remove cellular and bacterial contaminants and the resulting filtrate was used for viral DNA extraction using the QiAmp Ultra-Sensitive Virus Kit (Qiagen, Hilden, Germany) and whole genome amplification using the TruePrime WGA Kit (Syngen Biotechnology, Inc, Taipei City, Taiwan), each according to the manufacturer’s protocol. The resulting DNA was sheared to 600 bp prior to library preparation using the NEBNext Ultra II Library preparation kit (New England Biolabs, Ipswich, MA, USA) according to the manufacturer’s protocol. The libraries were then sequenced as 150 bp paired-end reads on the Illumina HiSeq 2500 platform (Illumina, Inc, San Diego, CA, USA). During that study (5), we identified one participant with a metagenome which consistently showed the presence of a feline papillomavirus and who also self-identified as being an owner of a domesticated house cat.

To further investigate this observation, 15 metagenome samples from that specific human individual were mapped to a *Papillomaviridae* reference database compiled from NCBI using BBMap v38.94 with a minimum match of 95% bp similarity and a k-value of 13 (6). Mapped reads were then processed using KHMER v2.0.0 (7) to remove singletons and overly-abundant reads using k-mer profiles. All filtered papillomavirus reads were then de-novo assembled using MEGAHIT v1.2.8 (8) and the subsequent assembly quality was assessed using QUAST v5.0.2 (9). Read coverage for the assembly was an average of 295.18 with a standard deviation of 46.61 reads per base. To quality check the metagenome assembly, the putative viral contigs were evaluated using CheckV v0.7.0 (10), and classified using nucleotide-based classification tools, such as Kraken2 v2.0.8-beta (11), Demovir (https://github.com/feargalr/Demovir), and Blastn (with a >10% query coverage cut-off) (12), as well as the protein-coding classification tool Kaiju v1.7 (13). All tools were run with default parameters unless otherwise specified.

A complete circular feline papillomavirus genome assembly was identified which showed 74% sequence similarity to feline papillomavirus type 2 (family *Papillomaviridae,* genus *dyothetapapillomavirus*). Genes of this single virus were annotated with PROKKA v1.14.5 (14) using the embedded viral annotation database and Cenote-Taker 2 v. 2.1 (15). This genome, which we identified as strain P20, was 8069 bp in length and exhibited a GC content of 54.38%. Six open reading frames were annotated, including the E6 Protein, E7 protein, E1 protein, E2 protein, Late Protein L2, and Major capsid protein L1 – which are shared among many animal papillomaviruses (1) (Figure 1; visualized with SnapGene, GSL Biotech LLC, San Diego, CA; https://snapgene.com). Using BLASTn, our genome assembly showing the greatest similarity (with an average of 70.3% shared amino acids) to a papillomavirus isolated from the skin of a domestic Maine Coon house cat in 2007 (GenBank accession NC038520).

**Figure 1.**
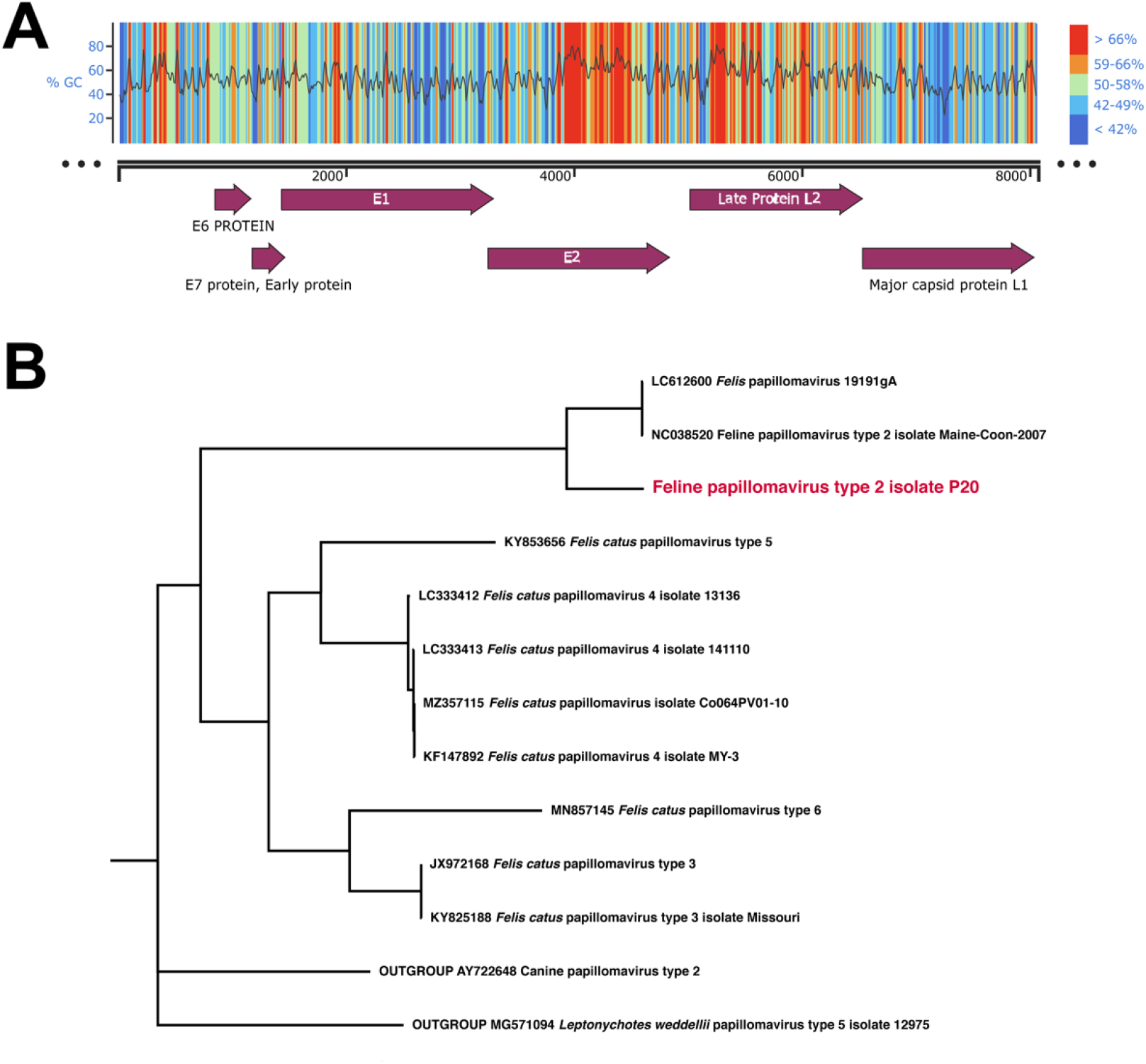
**A.** Linear genome map, 8069 bp in length, of Feline papillomavirus type 2 like virus. Top portion of the figure shows %GC content for regions of the circular genome. Higher GC content are shown in warmer colors (red) and lower GC content are shown in cooler colors (blue). The location of the six gene regions in the genome are shown below which consist of open reading frame annotations (purple arrows) of the E6 Protein, E7 protein, E1 protein, E2 protein, Late Protein L2, and Major capsid protein L1. **B.** Phylogenetic tree of the placement of the feline type 2 papillomavirus identified in this study along with nine other feline papillomaviruses. The tree was rooted with Canine papillomavirus type 2 and *Leptonychotes weddelii* (Wedell Seal) papilloma virus type 5.

To determine the phylogenetic placement of strain P20 identified here, we downloaded all feline papilloma genomes available on NCBI (along with two outgroups from closely related hosts – domestic dog and Weddell Seal) then aligned the six shared annotated genes using MUSCLE v3.8.1551 (16). After removal of uninformative amino acid regions using GBLOCKS v0.91b (17), a phylogenetic tree (Figure 1) was generated using IQTREE v1.6.12 (18) under the LG+F+I+G4 substitution model optimally determined by MODEL-FINDER v1.5.4 (19). This phylogeny, visualized with FIGTREE v1.4.4 (https://github.com/rambaut/figtree), placed our genome in proximity to other feline papilloma type 2 accessions.

## DATA AVAILABILITY

All raw sequencing data has been deposited in the NCBI Short Read Archive (SRA) under the accession code PRJNA754140. Within that project, the 15 sequencing files used in this study were the following: SRS9770471, SRS9770472, SRS9770476, SRS9770495, SRS9770500, SRS9770501, SRS9770587, SRS9770588, SRS9770589, SRS9770614, SRS9770616, SRS9770617, SRS9770642, SRS9770643, SRS9770644. The complete feline papillomavirus P20 genome has been submitted to NCBI Genbank with the accession number of OL310516. All metadata, sequences, annotation files, and scripts used here are publicly available and archived at: https://github.com/HerrLab/Graham_2021_feline_papilloma_P20.

## ACKNOWLEDGMENTS

This work was completed using the Holland Computing Center of the University of Nebraska, which receives support from the Nebraska Research Initiative.

We thank all the anonymous participants for their contribution to this study. We also thank members of the Fernando Lab who assisted with advice on laboratory methods.

